# A computational model that explores the effect of environmental geometries on grid cell representations

**DOI:** 10.1101/152066

**Authors:** Samyukta Jayakumar, Rukhmani Narayanamurthy, Reshma Ramesh, Karthik Soman, Vignesh Muralidharan, V. Srinivasa Chakravarthy

**Affiliations:** Dept. of Biotechnology, Rajalakshmi Engineering College, Chennai, Tamil Nadu, India; Bhupat and Jyoti Mehta School of Biosciences, Department of Biotechnology, Indian Institute of Technology Madras, Chennai, Tamilnadu, India-600036.

## Abstract

Grid cells are a special class of spatial cells found in the medial entorhinal cortex (MEC) characterized by their strikingly regular hexagonal firing fields. This spatially periodic firing pattern was originally considered to be invariant to the geometric properties of the environment. However, this notion was contested by examining the grid cell periodicity in environments with different polarity (Krupic *et al* 2015) and in connected environments (Carpenter *et al* 2015). Aforementioned experimental results demonstrated the dependence of grid cell activity on environmental geometry. Analysis of grid cell periodicity on practically infinite variations of environmental geometry imposes a limitation on the experimental study. Hence we analyze the grid cell periodicity from a computational point of view using a model that was successful in generating a wide range of spatial cells, including grid cells, place cells, head direction cells and border cells. We simulated the model in four types of environmental geometries such as: 1) connected environments, 2) convex shapes, 3) concave shapes and 4) regular polygons with varying number of sides. Simulation results point to a greater function for grid cells than what was believed hitherto. Grid cells in the model code not just for local position but also for more global information like the shape of the environment. The proposed model is interesting not only because it was able to capture the aforementioned experimental results but, more importantly, it was able to make many important predictions on the effect of the environmental geometry on the grid cell periodicity.

## Introduction

Spatial navigation is essential for the survival of a mobile organism. Entorhinal cortex (EC), an important sub-cortical area that forms input to the hippocampus, was reported to have neurons known as grid cells which fire when the animal is at points that have a spatially periodic structure(Hafting, Fyhn et al. 2005). Since the periodicity encountered is often hexagonal, these cells are further known as hexagonal grid cells(Hafting, Fyhn et al. 2005). Albeit grid cells were initially discovered in rats(Hafting, Fyhn et al. 2005), these cells have also been reported in mice(McHugh, Blum et al. 1996; Rotenberg, Mayford et al. 1996; Fyhn, Hafting et al. 2008), bats(Ulanovsky and Moss 2007; Yartsev, Witter et al. 2011), monkeys(Ono, Nakamura et al. 1993; Rolls and O'Mara 1995; Rolls, Robertson et al. 1997; Killian, Jutras et al. 2012) and humans(Ekstrom, Kahana et al. 2003; Jacobs, Weidemann et al. 2013; Moser, Roudi et al. 2014). Experimental studies in human adults who are at genetic risk for Alzheimer’s disease have reported that the neural degeneration originates in the EC, with the loss of grid cell representations causing further impairment of spatial navigation performance of the patient (Kunz, Schröder et al. 2015).

Preliminary studies on the effects of environmental geometry on spatial cells such as place cells (Jeffery and Burgess12 2006) and grid cells have been conducted. Place cells are critical for coding the animal’s position in space. They fire when the animal is situated in a particular space of the environment known as its firing field (O'Keefe and Dostrovsky 1971). Remapping of place cells occurred when sufficient changes to the geometry (Lever, Wills et al. 2002), color (Bostock, Muller et al. 1991) or odor (Anderson and Jeffery 2003) of the environment were made. Grid cells are equally crucial for spatial navigation by path integration i.e., tracking position by integrating self-motion even without the presence of external sensory landmarks(Hafting, Fyhn et al. 2005; Fuhs and Touretzky 2006; McNaughton, Battaglia et al. 2006; Burgess, Barry et al. 2007; Hasselmo, Giocomo et al. 2007; Moser, Roudi et al. 2014; Bush, Barry et al. 2015). Grid cells have been proposed to have a role in computing directional vector between the start and goal location (which was termed as vector navigation) that further aids the animal in reaching its goal location(Bush, Barry et al. 2015). The variation of the grid scale across the dorsal to ventral medial entorhinal cortex (MEC) axis(Brun, Solstad et al. 2008; Stensola, Stensola et al. 2012), acts like a ruler with different resolutions to measure the distance traversed by the animal from its starting location. The aforementioned features of the grid cells help the animal to navigate the environment efficiently.

MEC conveys spatial information into the hippocampus (Barnes, McNaughton et al. 1990; Quirk, Muller et al. 1992;Fyhn, Molden et al. 2004). It is believed that the dynamic representation of the spatial location of an animal is created and updated by the MEC and the grid cells are a proof of it(Savelli, Yoganarasimha et al. 2008). Hargreaves et al(2005) initially recorded MEC grid cells from an environment (Hargreaves, Rao et al. 2005). Since the size of the environment was relatively small, the neurons did not show obvious grid like firing patterns. It was very ambiguous in prior studies whether all spatially modulated cells in the MEC were variants of the grid cells or whether a subset resembled the place cells of the hippocampus (Savelli, Yoganarasimha et al. 2008). Savelli *et al* (2008) conducted an experiment where the rats were allowed to forage a small box which was placed inside a larger box. After sometime, the small box was removed from the large box without removing the rats and now the rats foraged the larger box for the rest of the experiment. It was observed that some cells that showed place cell like response in the small box, showed grid cell like response in the larger box and the cells showing boundary cell like response showed no change upon the removal of the small box. The aforementioned experiments gave two inferences: Firstly, it put forth that there were two major classes of spatial neurons, the grid cells and the boundary cells. The boundary cells may therefore be binding the grid cell firing to the boundaries of the environment. Secondly, the experiment strongly demonstrated that the spatial firing of the MEC cells were strongly influenced by the boundaries of the environment, in the sense that representation of the MEC cells changed predominantly when the local cues of the environment were altered. In this paper, we are addressing the second inference from a purely computational point of view.

The minimal remapping of the grid fields across the environment made grid cells to be the universal metric for navigation (Hafting, Fyhn et al. 2005). But this feature of grid field invariance across the environment was contested by the experiment conducted by O’Keefe(Krupic, Bauza et al. 2015)wherein rats were allowed to forage inside differently shaped environments such as circle, square, hexagon and trapezoid. Analysis of grid cell activity in each environment revealed that the hexagonal grid field symmetry was affected by the symmetries of the environmental shape. Circle, the most symmetric environment, had a regular hexagonal firing field. As the number of axes of symmetry dropped, the regular hexagonal firing field started to transform into a skewed hexagonal field. This experimental study pointed out that grid cell firing fields were not invariant with respect to the environment but exhibit a definite dependence on the geometry of the environment.

Another interesting experimental study(Carpenter, Manson et al. 2015)considered how grid cells responded when the animal foraged inside similar environments connected by a corridor. A key result of the study was that initially the grid fields in each room had a high spatial correlation between them; as the time progressed, this correlation decreased and the grid fields in the two environments became a continuum, forming a global representation of the connected pair of environments. This study revealed a new face of the grid cell coding, whereby the periodic firing fields of the grid cell could rearrange among themselves to reflect the global shape of connectivity of the environment.

Most of the experimental studies on grid cells were performed on either square or circular environments, and have not explored the rich possibilities of environmental geometries. Apart from the study of Krupic*et al* (2015) (Krupic, Bauza et al. 2015),to the best of our knowledge, no experimental studies have been conducted on grid cells under varying environmental geometries. Theoretical studies have been made on grid cell coding in non-Euclidean space(Urdapilleta, Troiani et al. 2015)which came up with the prediction of variation in the hexagonal pattern of firing field to heptagonal pattern with the change from Euclidean to non-Euclidean space. But the problem of studying grid cell coding as a function of practically infinite variations of the environmental geometry poses a Himalayan challenge to spatial cell researchers. Hence, we propose to classify environmental geometries into the following four broad categories and study the emergent grid fields using computational modeling.

i. Connected environments
ii. Convex shaped environments
iii. Concave shaped environments
iv. Regular polygon environments with varying number of sides

The aforementioned studies, from a pure computational point of view, would result in a better understanding of the spatial encoding inside the brain done by the grid cells. We show that our simulations not only explain and confirm earlier studies, but also make a number of testable predictions verifiable by experiment.

## Methods

To achieve the goal of studying the effect of environmental geometry on grid cell coding, we used the model described in(Soman, Muralidharan et al. 2016) with slight modification (Fig. 1) as explained below.

**Fig. 1:**
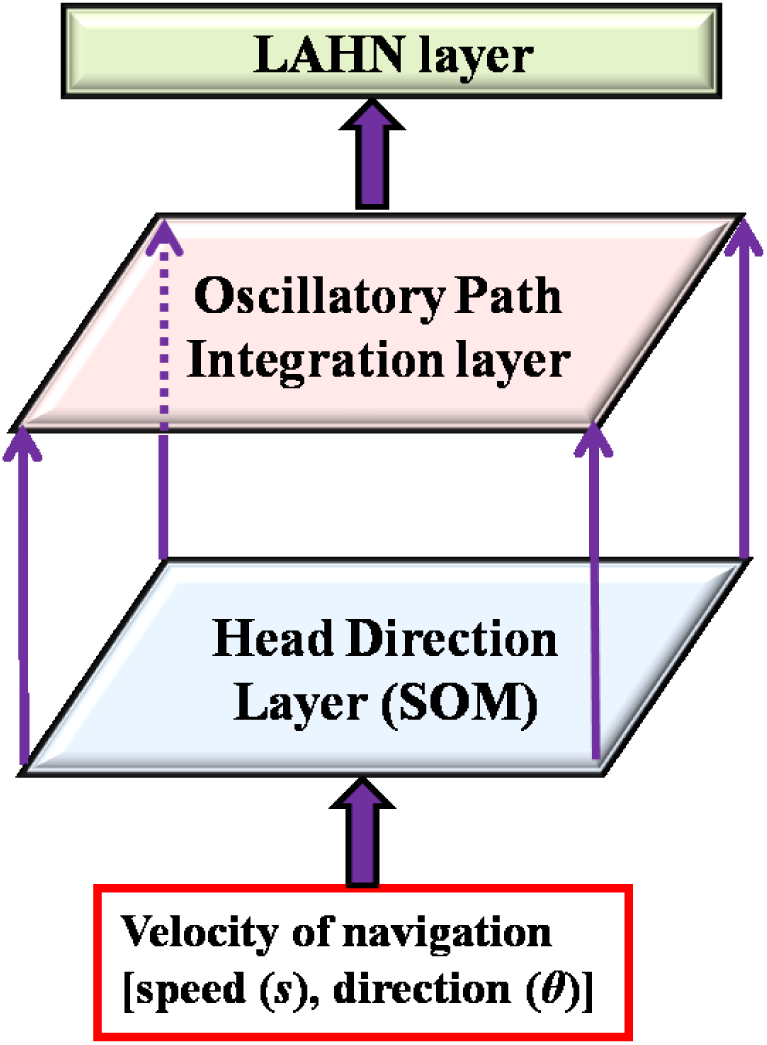
Model architecture. Arrows show the direction of flow of information from one hierarchy to the next.

The model basically has three stages as described below.

a. *Direction encoding stage* This Head Direction (HD) stage forms a neural representation for the direction in which the animal moves. In our earlier model, the direction features were(Soman, Muralidharan et al. 2016) extracted from the limb oscillations of the animal that made the model more biologically plausible. But here we simplified this stage by directly supplying velocity inputs to the model so that additional constraints on the curvature of the trajectory could be circumvented. To obtain a directional map, a Self Organizing Map (SOM) was trained(Kohonen 1982) using two dimensional inputs from a unit circle. The response equation of the SOM neuron is given as:
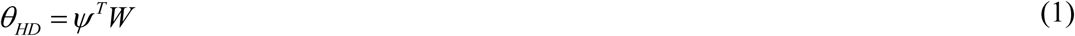 *ψ* is the two dimensional input given to the SOM such that ψ = [cos(θ) sin(θ)] where θ is the actual direction of navigation *W* is the afferent weight matrix of the SOM, where each weight vectors are normalized.
b. *Oscillatory Path Integration (PI) stage* This stage consists of a two dimensional array of Hopf phase oscillators, which has one-to-one connections with the HD layer. The directional input from eqn. (1) is fed to the phase dynamics of the oscillator so that each component of the positional information is encoded as the phase of each oscillator. The dynamics of phase oscillator is given as
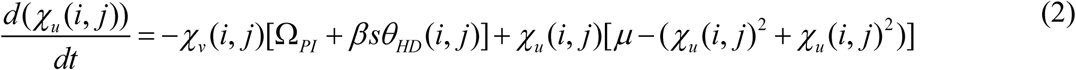

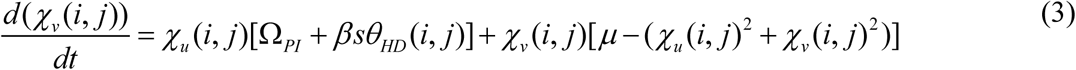 χ is the state variable of the PI oscillator. *β is* the spatial scale parameter. *s* is the speed of the navigation such that*s* = ||X(t)-X(t-1)|| where X is the position vector of the animal. *μ* is the parameter that controls the limit cycle behavior of the oscillator. Here *μ* is taken as 1.
c. *Lateral Anti Hebbian Network (LAHN) stage* LAHN is an unsupervised neural network(Földiák and Fdilr 1989) that extracts the high variance features from the input. The network has Hebbian afferent and anti-Hebbian lateral connections. The response of the network is given by the following equation.
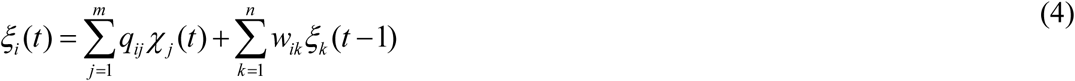 *q* is the afferent weight connections and *w* is the lateral weight connections. *ξ* is the response of the network. The afferent connections are updated by Hebbian rule and the lateral connections are updated by Anti-Hebbian rule as given below.
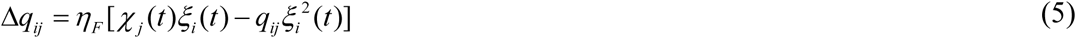

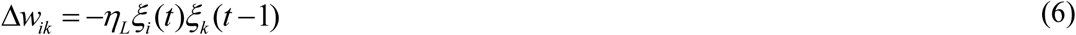 *η_F_* and *η_L_* are the forward and lateral learning rates respectively. The network is trained until a statistical condition such that the ratio of the mutual information of the network to the mutual information of Principal Component Analysis (PCA) on the input data reaches a pre-defined threshold(Oja 1982; Sanger 1989).

### Quantification of gridness

Although, once trained, the LAHN layer in the above model exhibits a variety of spatial cells, we primarily focused on the hexagonal grid cells to compare with the experimental results. Hexagonal gridness was quantified by a gridness score value(Hafting, Fyhn et al. 2005) computed from the autocorrelation map, obtained using the following equation.
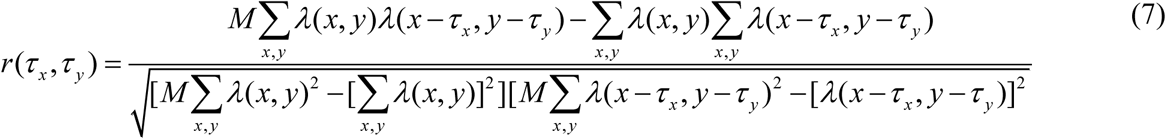

r is the autocorrelation map.

*λ(x,y)* is the firing rate at (x,y) location of the rate map.

*M* is the total number of pixels in the rate map.

*τ_x_* and *τ_y_* corresponds to x and y coordinate spatial lags.

Hexagonal Gridness Score (HGS)was computed as given below.
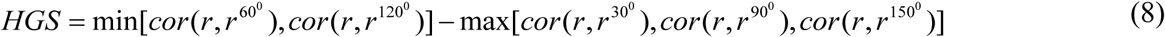

### Generation of Regular Polygons with varying number of sides

The polygons were constructed using a unit circle with centre at (0,0). The circle was then sectored into equal angular separation based on the number of sides given as the input.

The positional coordinates of the points on the unit circle that form the polygon are given by [X_i_Y_i_] = [cos(2π/n)sin(2π/n)]

Angle separation = 2π/n; n = number of sides

The X and Y coordinates were then connected to generate the regular polygon (Fig. 12).

### Generation of connected environments

The connected environment used for our study had the same boundary conditions used in the experiment (Carpenter, Manson et al. 2015). The two compartment (for e.g. square-square) connected via a rectangular corridor was constructed by joining the corner coordinates (Fig.2).For connected environments with varying distances between the two compartments, we introduced a distance parameter ‘*d*’. The displacement between the two squares is parallel to one side of each of the two square environments (Fig. 6 A, B and C).

**Fig.2:**
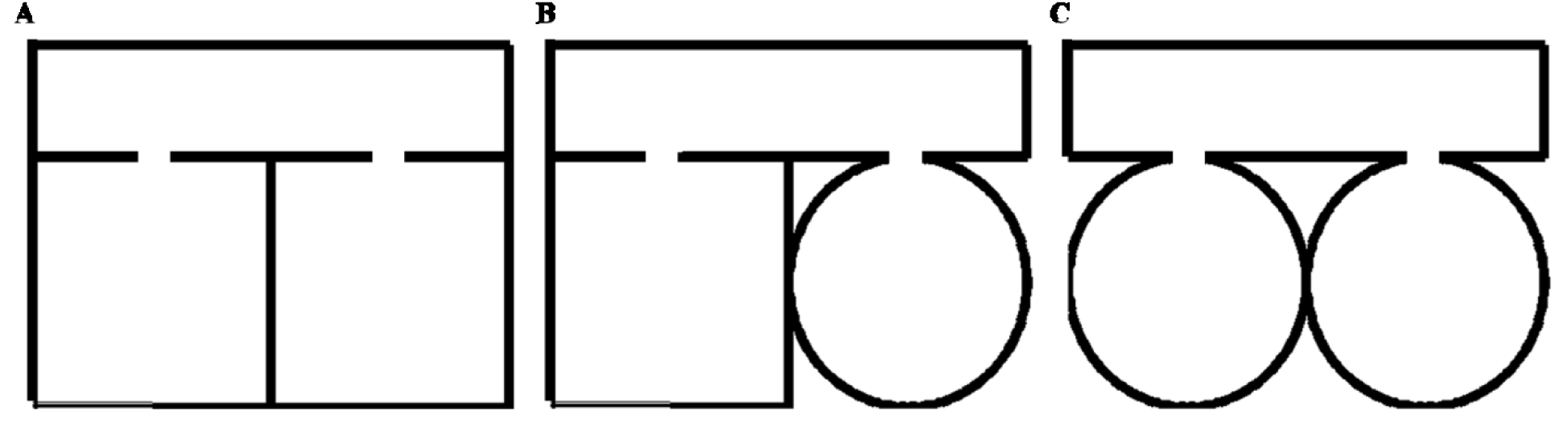
Boundaries of (A) square–square, (B) square–circle and (C) circle–circle connected Environment.

### Generation of concave boundaries

In this category, we consider the annulus, horseshoe and S-shape as instances of concave shapes. To construct an annulus shape, two circles were generated separately (of different radius) and then concatenated together to form a concentric circle (Fig.8A). In the case of a horseshoe (Fig. 8B), the starting and ending points of the shape were given as inputs along with coordinates obtained by the following equation.

[X_1_,Y_1_] = [-r_1_cos(θ) r_1_sin(θ)]; corresponds to the outer arc

[X_2_,Y_2_] = [r_2_cos(θ) r_2_sin(θ)]; corresponds to the inner arc

r_1_andr_2_ = radii of the outer and inner arcs respectively

The S-shaped boundary was generated by concatenating two horseshoe boundaries, with one of the horseshoes inverted to form the S- shape (Fig. 8C).

## Results

### I. Grid cell spatial coding in connected environments

We performed two different studies to understand the grid cell coding that emerges when the animal foraged environments connected by a narrow corridor. In the first study we manipulated the shapes of the connected environments and analyzed the grid fields. In the second study we fixed the shape but varied the distance between the connected environments.

#### a. Manipulating the shapes of the connected environments

We simulated connected environments with boundaries and corridor in the same dimension (the dimension of the square room was 1.8 x 1.8 units and that of the corridor was 0.8 unit)as used in the experimental study(Carpenter, Manson et al. 2015). We verified grid cell coding under three schemes such as square-square, square–circle and circle–circle as shown in Fig. 2. The virtual animal was allowed to forage the environment in these three cases. For each case, the model was trained and the resulting grid fields were analyzed as shown in Fig. 3.

**Fig.3:**
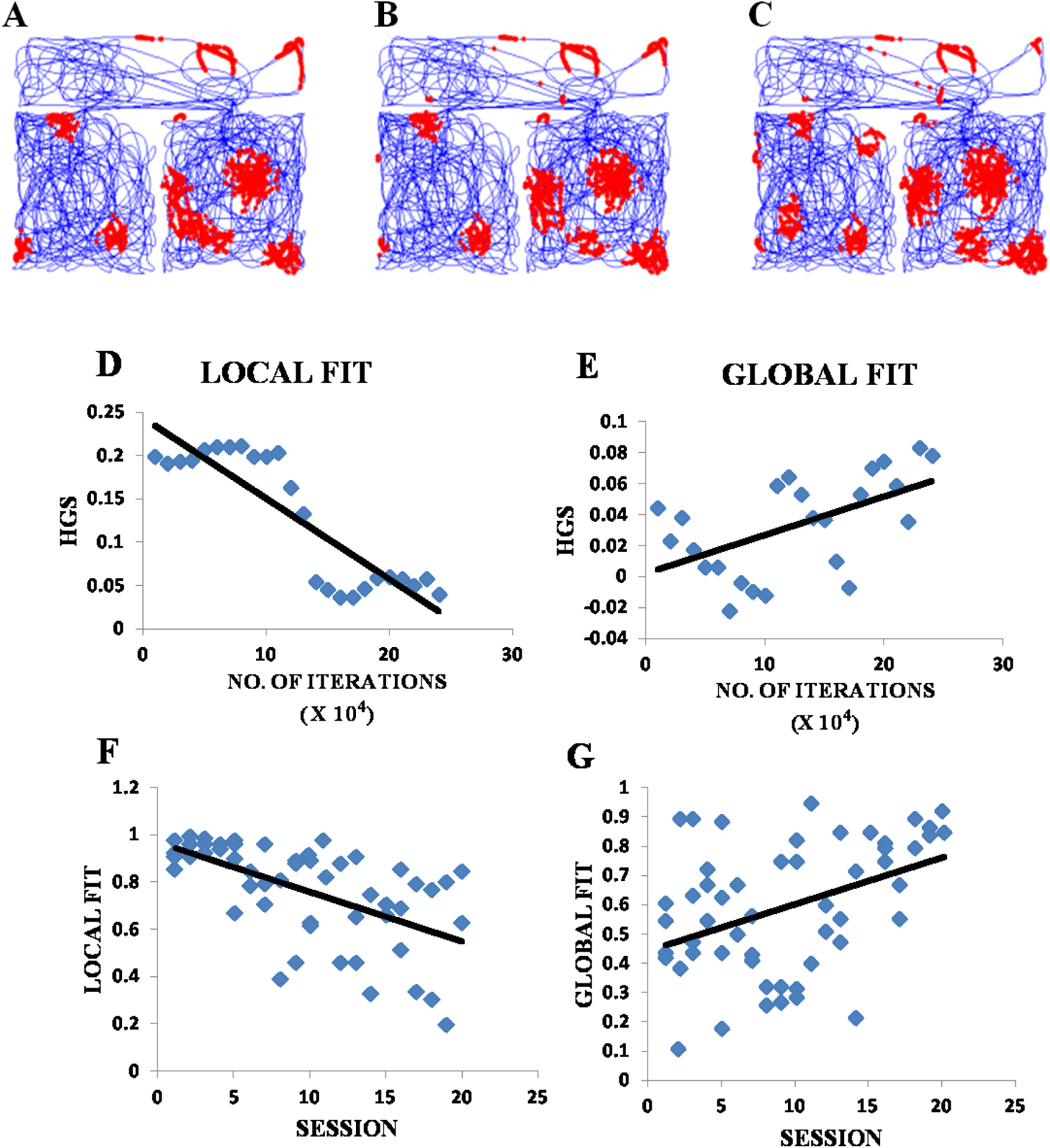
Grid cell firing in square-square connected Environment. A, B, C- represent the firing fields of grid cell neuron for square–square connected environment during different training iterations (beginning, middle and end respectively)of the model. D, E -Local and Global fit of the HGS values plotted against the no.of training iterations of model. F, G - Local and Global fit of the hexagonal fields in the square-square connected environment adapted from the experiment(Carpenter, Manson et al. 2015).

The global fit was computed by calculating the HGS values over the entire connected environment and the local fit was obtained by calculating the HGS values for the two square environments separately and averaging them. Global fit showed an increasing trend (Fig. 3E) with respect to the LAHN training time (Regression analysis: Global fit R^2^= 0.3826, p-value <0.05). Local fit showed a decreasing trend (Fig. 3D). (Regression analysis: Local fit R^2^=0.771, p-value < 0.001).

We carried out similar analysis for connected environments with different shapes such as square–circle. These boundaries were connected in the exact same manner as in the square–square case. In this case, the global fit showed a reverse trend (Fig. 4E) compared to the square–square case (Regression analysis: Global fit R^2^= 0.7042, p-value < 0.001) and the local fit showed an increasing trend (Fig. 4D). (Regression analysis: Local fit R^2^=0.8152, p-value< 0.001)

**Fig.4:**
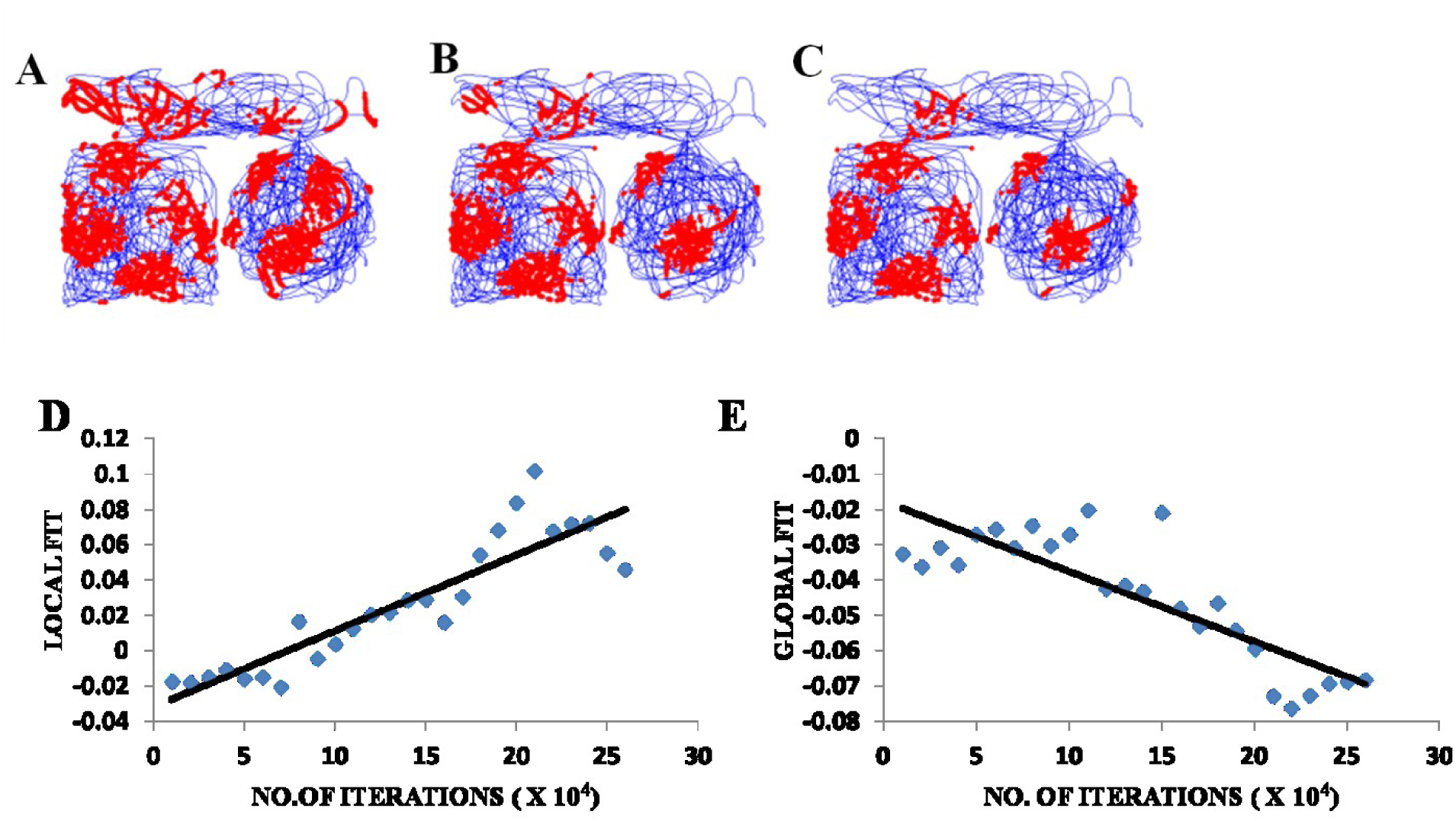
Grid cell firing in square-circle connected environment. A,B, C- represent the firing fields of the grid cell in the LAHN for square–circle connected environment at different training iteration (beginning, middle and end respectively) of the LAHN. D, E - Local and Global fit of the HGS values plotted against the no. of training iterations of LAHN.

We then connected circle-circle boundaries in the exact same manner as the ones before. Similar analysis was carried out and the global fit showed an increasing trend (Fig. 5E) (Regression analysis: Global fit R^2^= 0.4833, p-value < 0.001) and the local fit showed a decreasing trend (Fig. 5D) (Regression analysis: Local fit R^2^=0.4071, p-value < 0.001). These trends were similar to the square-square case.

**Fig.5.**
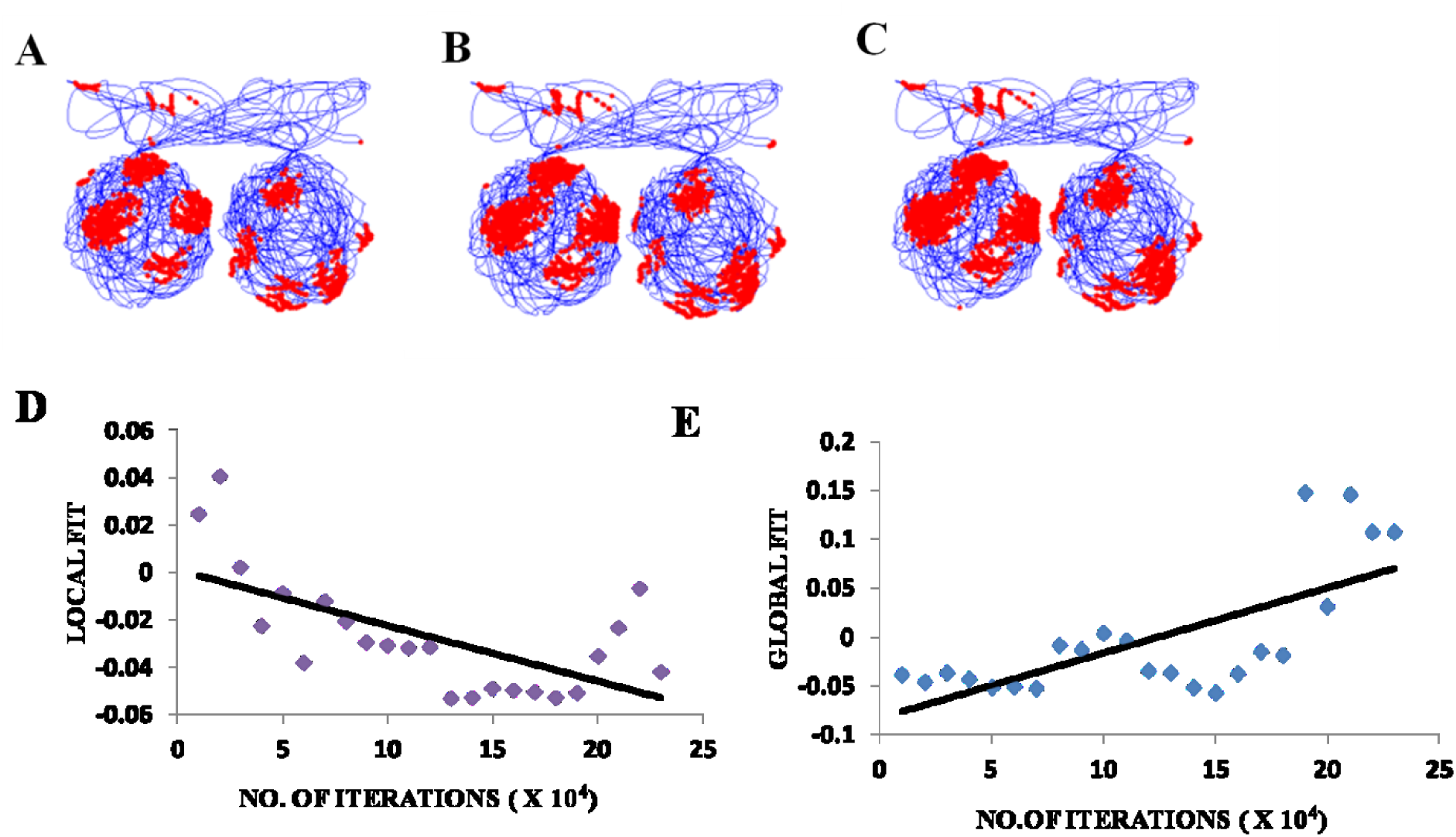
Grid cell firing in circle-circle connected Environment. A,B,C- represent the firing fields of the grid cell in the LAHN for circle–circle connected environment at different training iteration (beginning, middle and end respectively) of the LAHN. D, E –Local and Global fit of the HGS values plotted against the no. of training iterations of LAHN.

#### b. Manipulating the distance between the connected environments

Experimental reports proved that grid cells were capable of forming coherent global representations in a connected environment (Carpenter, Manson et al. 2015). Our next objective was to examine whether this globally representative property of grid firing is retained with increasing distance between the connected environments. To this end, we took square-square connected environment and varied the distance between the compartments. To examine whether this globally representative property of grid firing is retained even when the two compartments are away from each other, we varied the distance (d) ranging from 0.1 to 1 unit, in increments of 0.1. The boundary conditions of the compartments and the corridor were set in the same ratio as in the experiment(Carpenter, Manson et al. 2015). The agent was allowed to forage the environment for a period of 10 sessions. Each session consisted of five trips and for each session the distance between the two compartments was increased by a value of 0.1.

HGS values (HGS_final_) were computed at the time of convergence of LAHN and calculated the global and local fits. These scores were taken into account owing to the fact that convergence corresponds to the completion of the training session of the weights. The global fit of HGS values over the distances showed a decreasing trend (Fig. 6E) (Regression analysis: Global fit R^2^= 0.4585, p-value < 0.05) whereas the values of the local fit showed an increasing trend (Fig. 6D)(Regression analysis: Local fit R^2^=0.4142, p-value<0.05).

**Fig.6:**
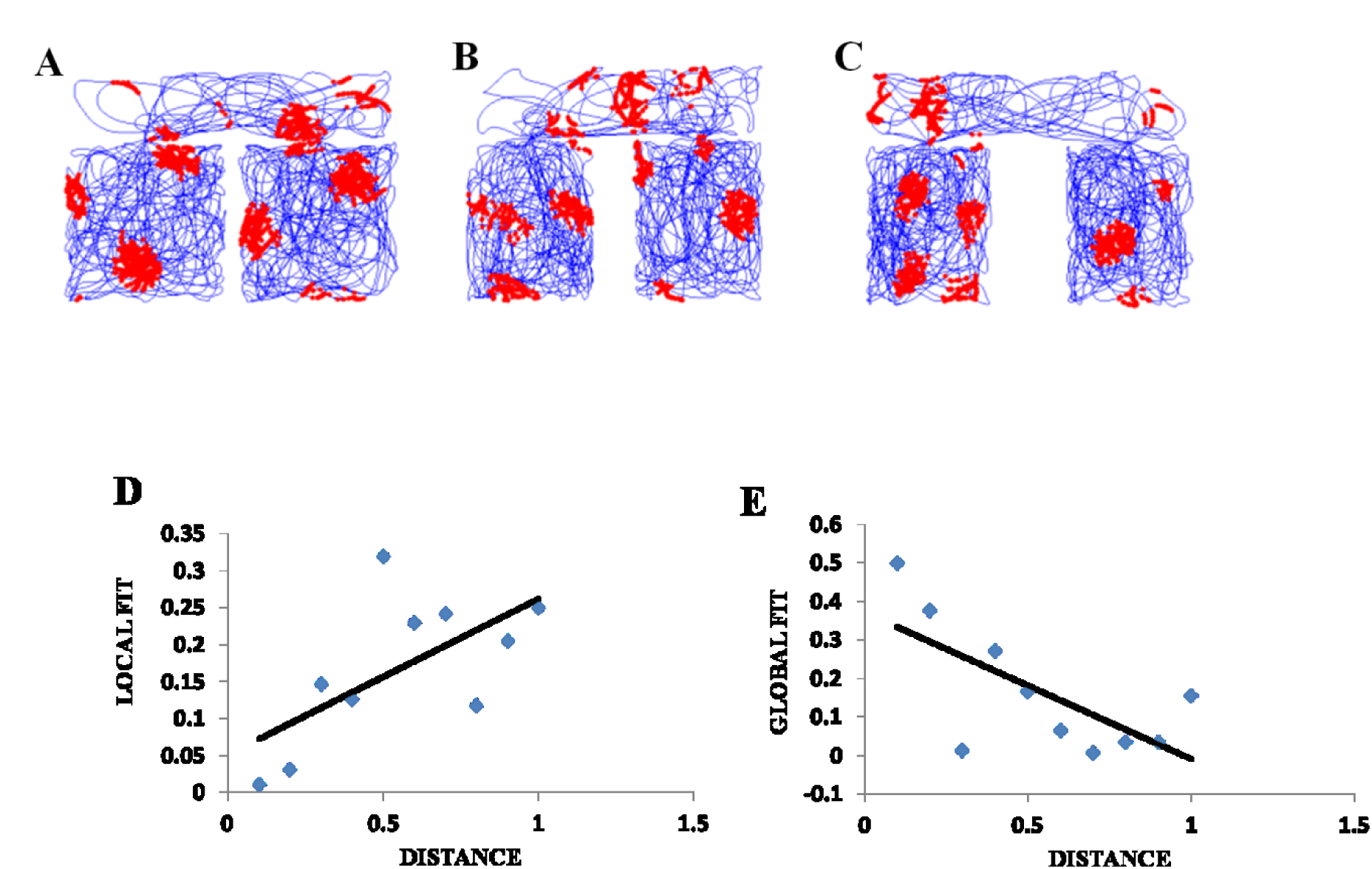
Grid cell firing in square–square Connected Environment with increasing distances. A,B,C – firing field maps (corresponding to end iteration of LAHN training) of the grid cell with respect to distances (d)= 0.1, 0.5 and 1unit respectively.D, E–Local and Global fit of HGS _final_with respect to the distance between the compartments.

### II. Grid cell spatial coding in convex shaped environment

The influence of environmental geometry on grid cell symmetry was contested by Krupic *et al* (2015) (Krupic, Bauza et al. 2015)and the notion that grid cells can serve as a universal metric for navigation was challenged. We conducted a similar study using our computational model wherein we generated square and trapezoidal boundaries in the same dimensions as used in the experiment (Krupic, Bauza et al. 2015).The HGS values were calculated from the autocorrelation map. Both the square and trapezoid boundaries were divided into two halves of equal areas and analysis was performed to check the similarity between the two halves for both the boundaries. The HGS analysis was carried out independently for each side and then for the complete shapes in case of both trapezoid and square. Owing to the fact that the grid patterns were more elliptical in a trapezoid, we used an elliptical orbit to extract the local peaks surrounding the central peak in order to calculate the gridness. The average HGS for the left side of the trapezoid was less when compared to its right (left= 0.050763, right = 0.14143; Fig.7D) and was more or less equal for both sides of the square (left= 0.201337, right= 0.20981, Fig.7C).

**Fig.7.**
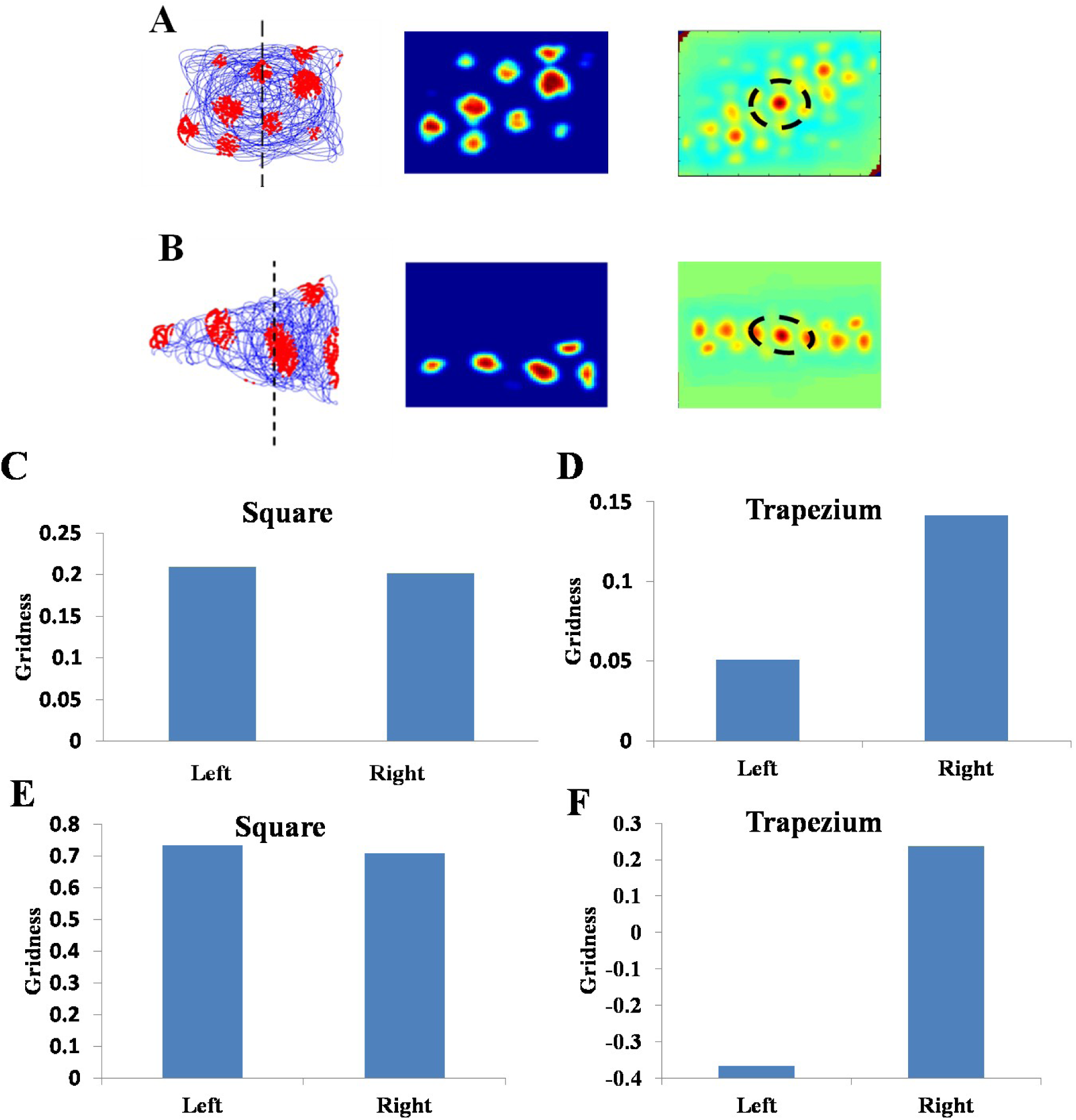
Grid cell firing in square and trapezoid enclosures. A, B – the firing field, firing rate and autocorrelation map of square and trapezoidal boundaries. The dashed vertical line in the firing field map divides the environment into two equal areas. Dashed circle and ellipse in the autocorrelation maps, show the best approximation of grid cell symmetry in square and trapezium respectively. C, D – Average HGS values plotted for the left and right orientation of the square and trapezium boundaries. E, F – Gridness of the left and right orientation of the Square and Trapezoid Boundaries adapted from the experimental study by (Krupic, Bauza et al. 2015).

### III. Grid cell spatial coding in concave shaped environment

A similar study of grid cell spatial coding was conducted using concave shaped environments like horseshoe, annulus and S shape as shown in Fig. 8.

**Fig.8:**
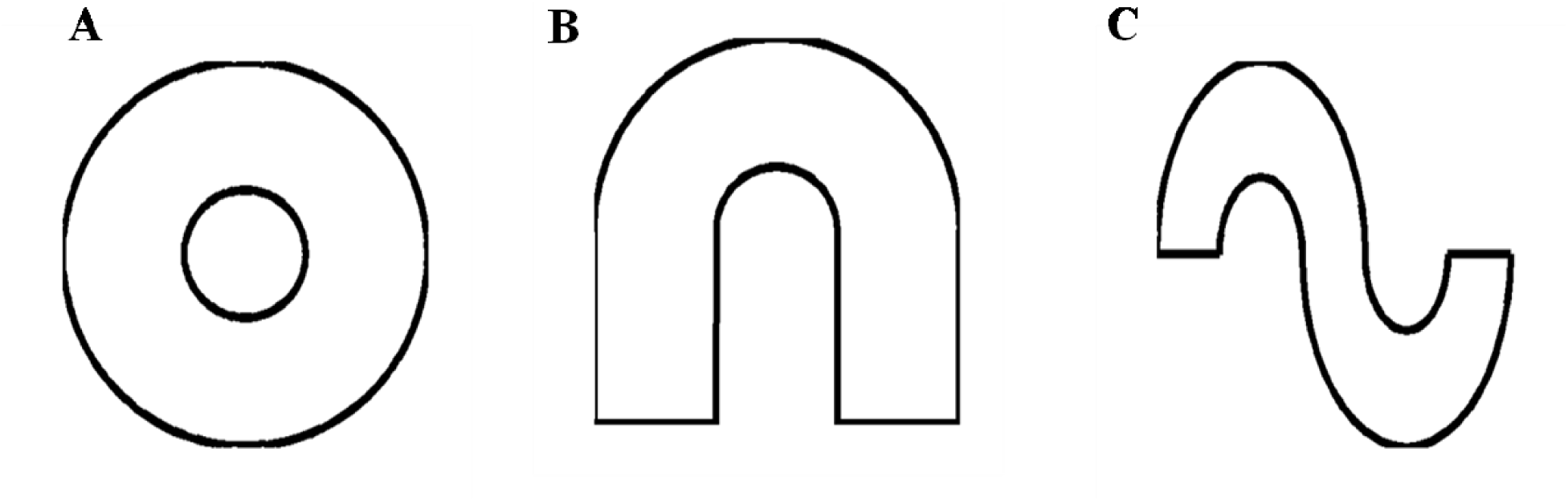
Simulated concave shaped environments such as (A) Annulus, (B) Horseshoe and (C) S – Shape respectively.

#### a. Horseshoe shaped environment

The inner radius (*r*) of the horseshoe was varied from 0 to 2 with a step size of 0.2. The horseshoe boundary with *r* = 0approximates a semicircle. The virtual agent was made to traverse the environment. Firing activity of the grid cells under various r values is shown as Fig. 9A-C. HGS values showed a decreasing trend (Fig. 9D) as the inner radius of the horseshoe was increased (Single factor ANOVA, p-value<0.001).

**Fig.9.**
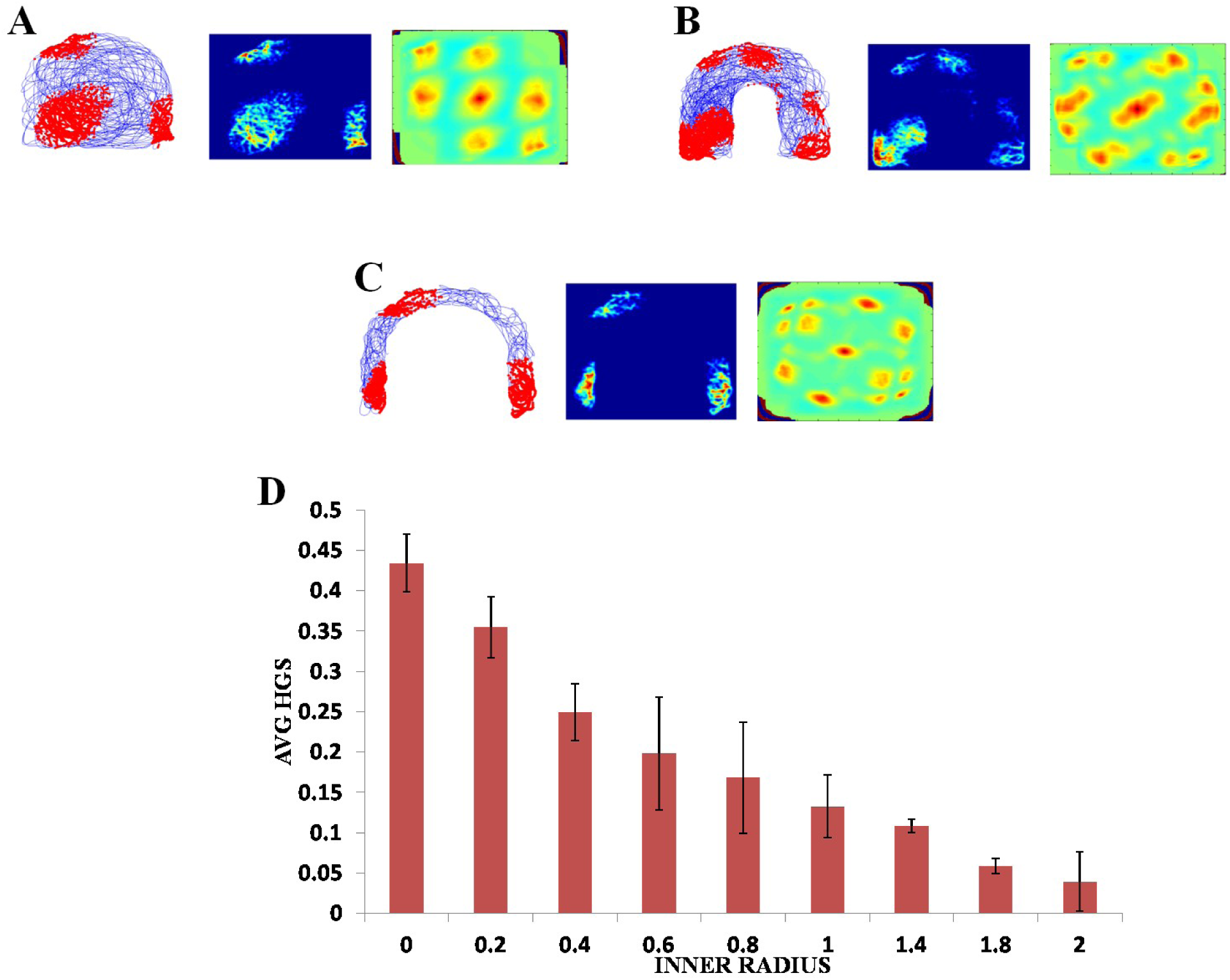
Grid cell responses in horseshoe shaped environment. A,B,C – Firing field, Firing rate and Auto Correlation maps of horseshoe boundary with varying inner radii of 0, 0.8 and 2respectively.D – Average HGS values vs inner radii of horseshoe shaped environment.

#### b. Annulus shaped environment

A similar analysis was performed with the second type of concave boundary i.e. annulus shaped environment. Annulus with inner radius 0 approximates to a circular boundary. The HGS values were computed and it was found to have the same decreasing trend as that of horseshoe as the inner radii of the annulus increased (Single factor ANOVA, p-value<0.001) (Fig. 10D).

**Fig. 10:**
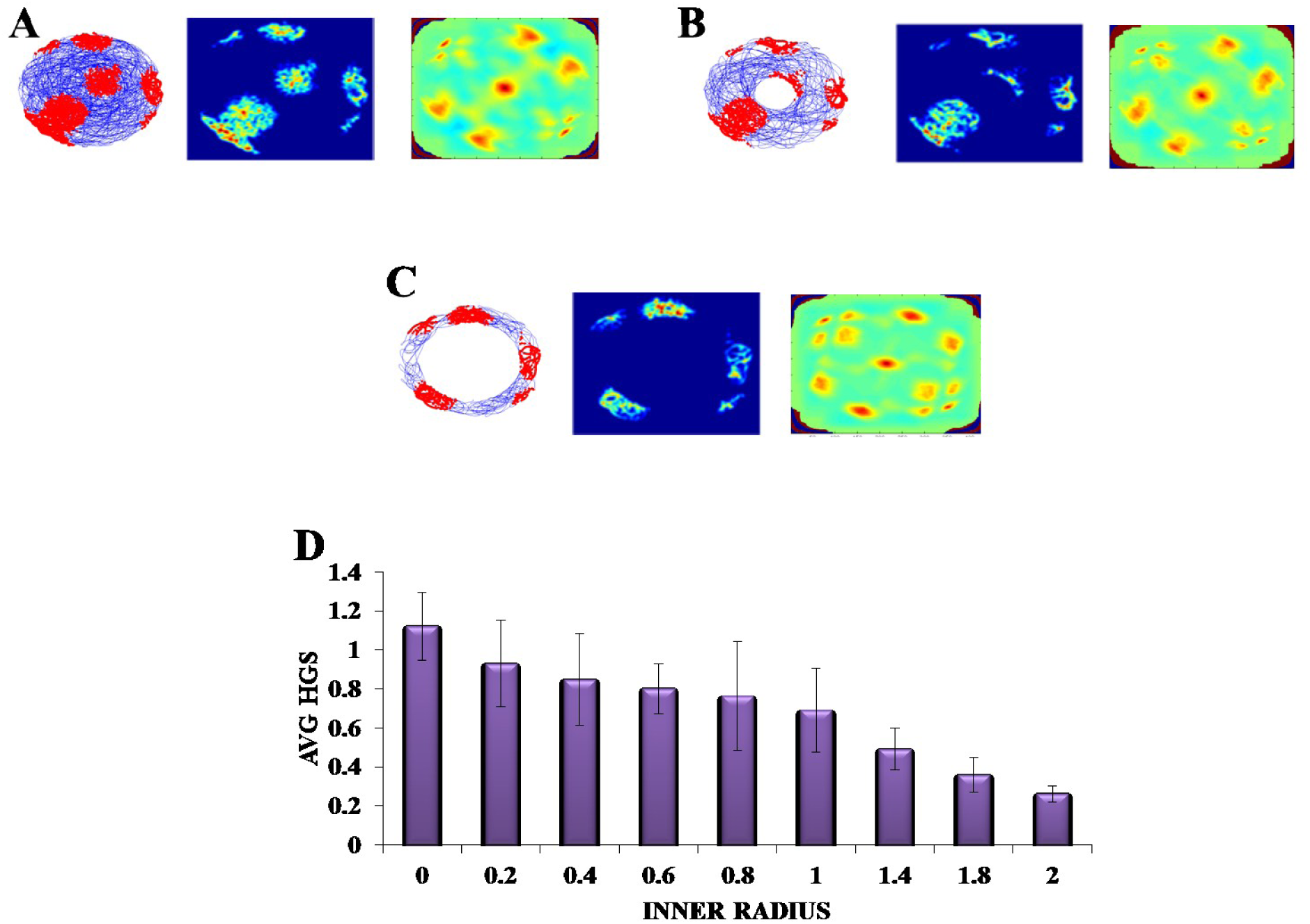
Grid cell response in annulus shaped environment. A,B,C – Firing field, Firing rate and Auto Correlation maps of grid activity in the annulus environment with varying inner radii of 0, 0.8 and 2 respectively. D – Average HGS values *vs* varying inner radii of annulus.

#### c. S shaped environment

In case of an environment like S shape, where two similar horseshoes were concatenated at a common end, it was found that as the inner radius of the S shape increased from 0 to 1 unit with a step size of 0.2, the HGS values showed a decreasing trend (Single factor ANOVA, p-value <0.001) (Fig. 11D).

**Fig. 11:**
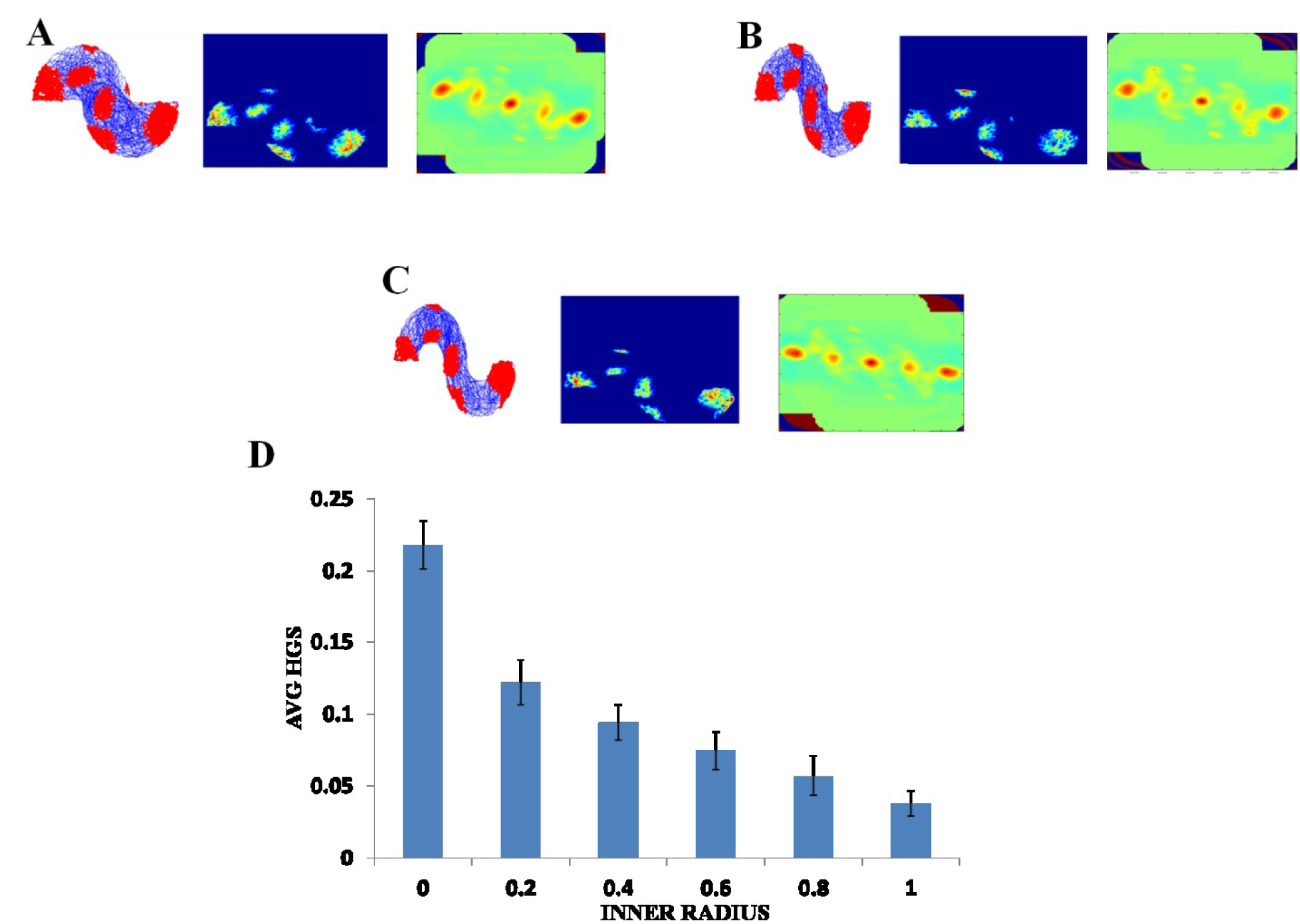
Grid cell response in S-Shape Boundary. A,B,C – Firing field, Firing rate and Auto Correlation maps of S shape boundary with varying inner radii of 0, 0.8 and 2 respectively. D – Average HGS values vs varying inner radii of S shape environment.

### IV. Grid cell spatial coding in regular polygons with varying number of sides

Most of the grid cell experimental recordings were carried out either in square or circular shaped environments (Krupic, Bauza et al. 2015). Here we address the problem of grid cell coding for environments in the shape of n-sided regular polygons. Specifically we consider the range of n from 3 to 10. This study will naturally include the square shape (four sided shape) and also will give an understanding on how the grid cell code will vary if the number of sides get increased and approximates a circle (a regular polygon with infinite sides). In other words, it is similar to study the influence of symmetry of the environment on the grid cell code. Fig. 12 shows the simulated environments used for this study.

**Fig. 12:**
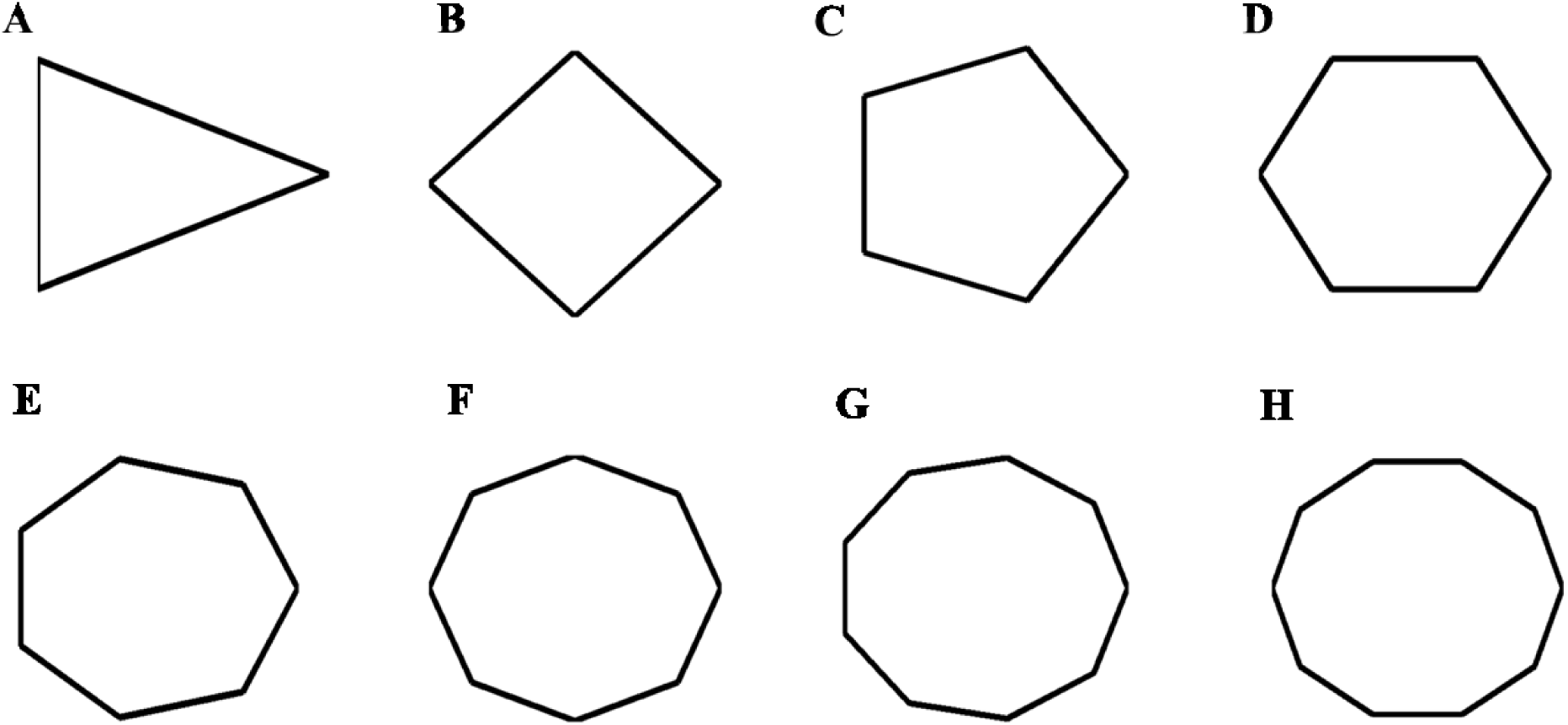
(A)-(H):Environmental shapes used for the analysis of grid cell coding with respect to the number of sides of the polygon; arranged in the order of triangle, square, pentagon, hexagon, heptagon, octagon, nonagon, decagon (from left to right).

The virtual animal was then made to forage inside these shapes and the resultant trajectory was given as input to the model. We computed the HGS values from the spatial autocorrelograms. It was observed that the HGS values showed an increasing trend with respect to the number of sides of the polygon (Single factor ANOVA, p-value <0.001) (Fig. 13I)

**Fig. 13:**
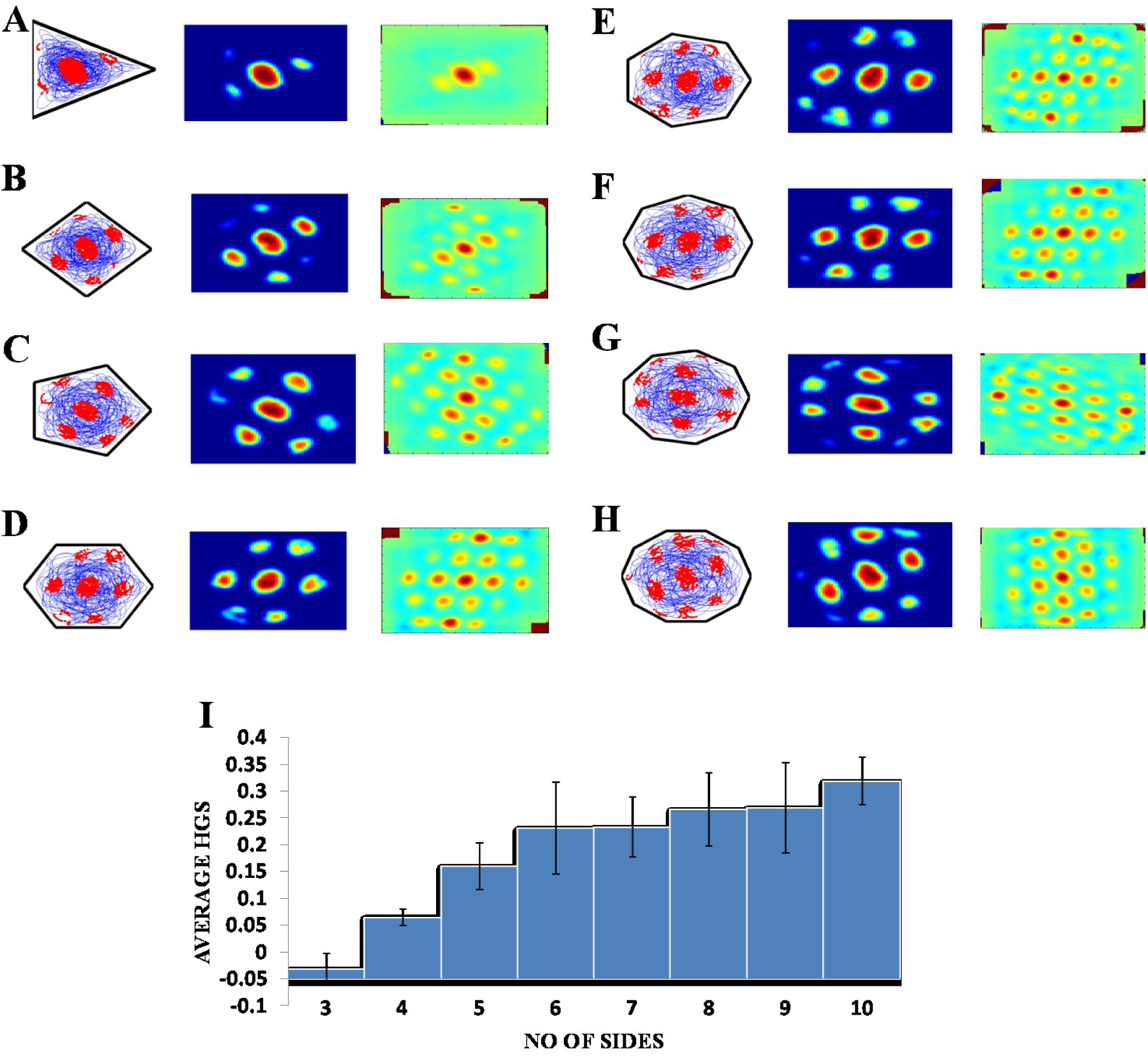
Grid cell firing in regular polygons.(A)-(H):Firing field, firing rate map and autocorrelogram (from left to right) of triangle, square, pentagon, hexagon, heptagon, octagon, nonagon and decagon shapes respectively. (I) the average HGS value of the grid field obtained for each polygon versus the number of sides of the polygon. The plot shows an increasing trend as the polarity of the environment decreases.

## Discussion

Grid cell firing fields, characterized by their hexagonal spatial periodicities, were considered to serve as a universal metric for spatial navigation. This notion was contested by some experimental studies (Krupic, Bauza et al. 2015) showing the dependence of grid cell coding on the environmental shape. But the experimental studies had limitations with regard to the grid cell recording under different environmental shapes. This forms the motivation of the present paper which seeks to study, using computational modeling, grid cell activity under different environmental geometries without any limitations that plague experimental efforts. Since we had to study the grid cell activity under various environmental conditions we systematically divided the simulations into four categories such as grid cell spatial coding in connected environments, convex shaped environments, concave shaped environments and finally variation of grid cell coding with respect to the number of sides of regular polygon-shaped environment.

In the connected environment study, we initially manipulated the shapes of the connected regions such as square-square, square-circle and circle-circle. HGS values were computed while the model was undergoing training. The interesting result was that in the case of similar shaped environments, such as square-square and circle-circle, the global HGS values (HGS value computed from the connected environment as a whole) showed an increasing trend and the local HGS values (average of the HGS values computed from each connected region separately) showed a decreasing trend with respect to the training time of the model. This means that as the animal gets more and more familiar with the environment (increasing the training time of the model), the grid fields start to realign themselves and form a continuum in the case of similar shaped connected environments. This result suggests that in addition to coding for the distance traversed by the animal, grid fields may also be implicitly coding for the global structure of the environment in which the animal navigates. In order to determine whether the global representation of the connected environments depends on the similarity between the two connected boundaries, we performed a similar analysis in a square-circle environment. But in this case a reverse result was obtained such that the simulation showed a negative trend in the global HGS and a positive trend in the local HGS value. This means that by analyzing the grid cell HGS variation, as the animal explores the environment, one may be able to infer whether the animal is exploring in a similar shaped environment or dissimilar shaped environment. This is a novel insight into the grid cell code purely from the computational point of view, and an easily testable prediction for future experiments.

After studying the grid cell coding scheme with respect to the shapes of the connected environments, we delved into the dependence of grid cell coding scheme on the distance between the connected environments. This study, along with the aforementioned study, is pertinent especially with regard to large scale navigation where the animal is not restricted to just one environment but shuttles between multiple environments of different shapes at different locations. Hence to get an insight on the grid cell coding with respect to the distance between the connected environments, in the simulation, we connected two similar shaped environments (square-square) and varied the distance between them. Our hypothesis was that since grid cells could code for the distance travelled by the animal(O'keefe and Burgess 2005) (due to its regularly periodic hexagonal firing field), the distan0063e between the connected environments should also be reflected in its activity. The analysis was the same as mentioned above. The variation in the global HGS values showed a decreasing trend and local HGS values showed an increasing which suggests that at greater distances, the coupling between the two environments is weakened. The grid cell representations generated treat the two environments as independent at larger distances. Hence at the outset, when the distance between the compartments was minimal, the representation was more global as opposed to local. As the distance between the two compartments increased, the grid cells seemed to lose their ability to form global representations and the firing was localized to their respective compartments. From the above simulations, the inference is that grid cell code may not be mere a distance code but it should be coding the entire structure of the environment and also probably the distance between the environments. The above methods can be easily extended to the case of connected environments with more than two components. We can consider a network of environments with complex spatial arrangements and connectivity. It would be interesting to study the evolution of local vs global organization of the grid fields in such system. In addition to the spatial arrangements of the environments, in such complex systems, even the frequency of visitation of that agent to individual components, may also determine the overall grid field organization. Such studies might pave way to the formulation of deep laws that govern the spatial encoding of brain in complex environments.

Oscillatory path integration stage of the model is vital to capture the aforementioned results. Here, the position is encoded as the phase of the oscillator (Eqn.2-3). If a grid cell is activated at one point in space in one case (for instance, in connected environments separated by distance d_1_) and not activated at the same point in the second case (distance d_2_), the reason must be that the afferent input to the grid cell from the oscillators is different (Eqn.4) in the two cases. Different configuration of the environment makes the oscillator to code for the same position at different phases of the oscillator. Also, since position is encoded as a periodic quantity at this oscillatory stage (as it does in oscillatory interference model (Burgess, Barry et al. 2007), this periodicity is reflected in the spatial firing fields of the grid cell in the LAHN.

The experimental study conducted by Carpenter et al(Carpenter, Manson et al. 2015) where the rat was made to forage between two similar square boxes connected via a corridor forms, the special case of the aforementioned modeling study where the shape is square-square and distance is zero. The modeling results concur with the experimental results (Fig.3). In the experimental case it was observed that initially the grid cell firing was controlled by the local cues, in the sense that the firing replicated between the two compartments. However, with further exploration, the similarities between the grid firing fields of the compartments decreased, suggesting that with increasing trials, global cues controlled the firing of grid cells. This trend was captured in the model also (Fig.3).Hence the aforementioned simulation and further analysis of the grid cell activity in the connected environments form a broad study which is easily testable in an experimental lab.

The objective of the next simulation study was to essentially capture the results of the experimental work by (Krupic, Bauza et al. 2015)where a rat was made to forage inside multiple boxes of different shapes such as circle, square, trapezium and hexagon. This study explained the permanent effect exerted by the environmental geometry on grid cell firing. The experiment demonstrated how the grid field symmetry was affected by the environmental geometry. To determine the impact of environmental characteristics on homogeneity and symmetry of grid patterns, the grid firing in two shapes such as square and trapezium was analyzed. It was found that in a highly polar environment like a trapezium there was a decrease in the regularity of the hexagon (reflected in the HGS score) and the pattern became highly elliptical across the entire enclosure. To estimate the regularity of grid patterns, the trapezoid and square was divided into two parts of equal area and the firing fields on both the sides were compared. The autocorrelation maps showed that there was a strong difference in local spatial structures between the two sides of a trapezoid unlike a square wherein they were highly similar. The gridness of left (narrower) side of the trapezoid was found to be low when compared to its right (broader) side. Also when the square and trapezoid boundaries were compared as a whole, the latter had a lower gridness. This was because when a trapezoid is divided into two, the left side resembled a triangle and the right side, a square.

In the simulation, we were able to capture the aforementioned results (Fig.7). We did the comparative study as mentioned above using our model i.e. between both sides of the trapezoid and square and between both the shapes as whole and obtained congruent results (Fig.7). From the firing rate map and autocorrelation map we were able to see that the left side of the trapezoid had less local spatial structure compared to its right side. In the case of a square, little difference was observed between its two halves.

Since the model successfully captured the experimental results(Krupic, Bauza et al. 2015) in the convex shaped environments such as square and trapezoid, we extended our study by varying the number of sides of a regular polygon. The aim of this study is to understand the effect of the number of degrees of freedom of the environment on the grid fields. Our simulations showed such that the HGS value showed an increasing trend with respect to the number of sides of the regular polygon (Fig. 13I). In other words, higher symmetry in the environment leads to higher HGS in the grid fields. Hence we predicted that the HGS score should be maximum for a circular environment (where the number of degrees of symmetry is infinite). It was also observed from the experiment(Krupic, Bauza et al. 2015), that the circular boundary is considered to be highly unpolarised when compared to all the other boundaries and hence showed high gridness scores.

Since the real world navigation occurs in environments with arbitrary shapes, we conducted the simulation studies on concave shapes too. We considered concave shapes such as horseshoe, annulus and S-shaped environments (Fig.8). As the inner radius of the horseshoe was increased, the hexagon formed in the auto correlation map appeared to lose its regularity. This was captured by the decreasing HGS values as shown in the graph (Fig.9D).This trend was observed for the other two concave boundaries as well, i.e. annulus and S-Shape (Fig.10D and 11D). It can be observed that, since the annulus with inner radius 0, approximates a circular boundary, its HGS value tends to be the highest when compared to the others. It was also observed that as the inner radii of the aforementioned concave environments was increased; the space available for the virtual animal to traverse shrunk. This reduced availability of space may be the reason behind the deviation from the hexagonal spatial coding of the grid cell, as reflected in the reduced HGS values across all the three concave shapes. This can be experimentally tested in many ways such as by implementing the environment similar to the one we simulated in this study (Fig.9-11) with less space given to the animal for exploration. Experimental studies have reported that medial septal cholinergic projections to the hippocampus modulate the exploratory behavior of the animal in such a way that increase concentration of acetylcholine in the septal-hippocampal projections reduced the exploratory behavior of the animal and vice versa (Lamprea, Cardenas et al. 2003). Therefore, a septal lesion study that affects the exploratory behavior of an animal can beyet another method for testing the aforementioned model results.

From the aforementioned simulation studies we propose relevant inferences regarding the grid cell coding. In all the boundary manipulation cases, we could see notable changes in the grid cell code which in turn was reflected in its firing fields and the corresponding HGS score. This means that grid cell activity may be coding not only for the distance metric but also for global information regarding the environmental geometry. If this is the case, then given the grid field activity, it should be possible to classify the environment in which the animal navigates and we consider this as the future work. In addition, we might also include the responses of all the LAHN neurons rather than considering only the grid cells for this study. If the LAHN activity as a whole gives a better classification of the environment, then it may point to the possibility that the animal may be relying upon the MEC activity as a whole (which is LAHN in the model) to comprehend its space, with the grid cells telling only part of the story. There is also a need to develop a strong mathematical framework that explains the relationship between the observed spatial coding and the geometry of the environment. One possible source of such a theory is the theory of contour integration from complex analysis, which allows us to specify the values of an analytic function at points inside a contour, based on the values of the function on the contour(Brown 2009). The validity of this insight needs to be ascertained by developing a suitable theory. We would also like to analyze the activity of other LAHN spatial cells with respect to change in the environmental geometry as the future work. Hence the proposed modeling study gives a new dimension to the grid cell coding with a good number of testable predictions.

